# Age and musical training effects on auditory short-term, long-term, and working memory

**DOI:** 10.1101/2022.08.31.506048

**Authors:** G. Fernández Rubio, E. R. Olsen, M. Klarlund, O. Mallon, F. Carlomagno, P. Vuust, M.L. Kringelbach, E. Brattico, L. Bonetti

## Abstract

Cognitive aging is characterized by a gradual decline in cognitive functioning. One of the most worrying deficits for older adults is a decreased capacity to memorize and remember new information. In this study, we assessed auditory short-term memory (STM), long-term memory (LTM), and working memory (WM) abilities of young and older adults using musical and numerical tasks. Additionally, we measured musical training and tested whether this capacity influences memory performance. Regarding STM, young adults scored higher than older adults when making same/different judgements of rhythmic sequences, but their performance was alike for melodic sequences. Higher levels of musical training were associated with enhanced STM capacity for melodic sequences. In relation to LTM, young adults outperformed older adults in identifying new musical sequences. Moreover, younger and older individuals with high musical training outperformed those with low musical training. No group differences were found in the recognition of previously memorized musical sequences. Finally, we found no group differences in WM capacity, although there was a non-significant tendency for young adults to outperform older adults. Overall, we found that aging differently affects several types of auditory memory and that, for certain musical memory tasks, a higher level of musical training provides significant advantages.

## Introduction

Normal aging is accompanied by a number of physical and psychological changes. Among them, cognitive aging, or the physiological and functional age-dependent changes in the brain, can become an important cause for concern in older adults.

Cognitive aging has been well documented in the scientific literature. Age-related declines in selective ^1, 2^ and sustained ^2^ attention are noticeable in many daily tasks (e.g., driving ^3^) and have been linked to overall white matter atrophy ^4^, reduced interhemispheric connectivity ^5^, and altered mechanisms in the frontoparietal network ^1, 6, 7^. Deficits in executive functioning also occur with advancing age, and have been described in rodents ^8^. Such deficits have been reported in cognitive flexibility ^9, 10^, planning ^11^, and inhibitory control ^6^. In addition, age-dependent deficits in processing speed can affect other cognitive domains ^12^.

One of the most investigated age-related cognitive decays is memory. Although memory functioning declines with advancing age, not all types of memory are equally affected ^13^. Semantic memory, which stores factual information and general knowledge, is relatively well preserved from early to late adulthood ^14, 15^. Similarly, procedural memory, or skill knowledge, is minimally affected by advancing age ^16, 17^. Conversely, episodic memory, which is long-term memory (LTM) of specific events, is highly age sensitive ^15, 18, 19^. So is working memory (WM), which is involved in the temporary storage and manipulation of information and has been suggested to decline due to decreased processing speed ^20^. Finally, short-term memory (STM), or the ability to maintain information after the sensory input has been removed, also declines with age ^21, 22^. Age-related memory deficits may be due to a number of neural factors. Buckner ^23^ concluded that disruption of the frontostriatal systems, medial temporal lobe and associated cortical networks accompanies deficits in attention, executive functioning, and long-term, declarative memory. These age-related factors all contribute to memory decline and vary greatly between individuals.

Within the field of cognitive aging, research on brain maintenance has quickly grown, particularly regarding compensatory recruitment. This term refers to the increased neural activation that is observed in elderly individuals when performing cognitive tasks. However, age-related under-recruitment of brain areas may also occur under certain circumstances ^24, 25^. In relation to memory, bilateral recruitment of frontal areas ^26, 27^ and posterior-to-anterior shifts ^28^ in activation patterns seem to facilitate encoding of episodic memories ^29^. Overactivation of prefrontal brain regions has also been observed in patients with Alzheimer’s disease during object location encoding and has been associated with compensatory mechanisms ^30^. Additionally, an age-related, load-dependent increase or decrease in the activity of the dorsolateral prefrontal cortex has been reported during WM tasks ^25, 31, 32^.

Related to compensatory recruitment, several mechanisms can mitigate cognitive decline in aging ^33, 34^. In the case of memory, educational level and IQ are associated with the use of effective memory strategies in older adults ^35^. Similarly, musical training has positive effects on cognitive functioning in the elderly. Seinfeld et al. ^36^ found that piano training improved performance on measures of visual scanning, motor ability, executive function, divided attention, and inhibitory control in a group of older adults. Additionally, Hanna-Pladdy and MacKay ^37^ showed that elderly musicians performed better than age-matched non-musicians in nonverbal memory, naming, and executive functioning tasks.

In this study, we focus on differences in auditory memory functioning between young and elderly adults and the effects of musical training on this cognitive domain. We wish to provide further information on this broad topic by comparing the performance of two age groups on several tasks that assess three types of auditory memory: STM, LTM, and WM. Some of the tasks involve music, so we also study how musical training level affects memory capacity. Following previous studies, we expect to observe a decline in elderly adults as compared to younger adults in all the investigated memory systems (STM, LTM, and WM). Furthermore, we hypothesize that the level of musical training will modulate this decline, especially in tasks comprising musical stimuli.

## Methods

### Participants

The sample consisted of 77 participants (43 females, 34 males) that were divided into two age groups. The young group comprised 37 participants (18 females, 19 males) aged 18 to 25 years old (mean age: 21.89 ± 2.05 years). The elderly group included 40 participants (25 females, 15 males) aged 60 to 81 years old (mean age: 67.50 ± 5.46 years). All participants were Danish and reported normal health and hearing.

The project was approved by the Institutional Review Board (IRB) of Aarhus University (case number: DNC-IRB-2021-012). The experimental procedures complied with the Declaration of Helsinki – Ethical Principles for Medical Research. Participants’ informed consent was obtained before starting the experiment.

### Materials and procedure

We tested participants’ individual STM, LTM, WM, and musical training levels in a quiet room at Aarhus University Hospital (Aarhus, Denmark).

The Musical Ear Test ^38^ (MET) was used to measure auditory STM abilities. The MET consists of 104 trials in which participants state whether two brief musical sequences are identical. The test is divided into two parts: Melody, which consists of 52 sets of melodic sequences, and Rhythm, comprising 52 sets of rhythmic sequences. Due to time constraints, we used a reduced version of the test consisting of 40 trials (20 sets of melodic sequences and 20 sets of rhythmic sequences) ^39, 40^. Scores ranged from 0 to 20 on each part.

To assess auditory LTM abilities, we used an old/new auditory recognition task. During the encoding part, participants were instructed to listen four times to a shortened version of Johan Sebastian Bach’s Prelude in C minor, BWV 847. Subsequently, during the recognition part, participants were presented with brief musical sequences that were extracted from the prelude (“old”) and with novel musical sequences (“new”) of the same length, and they were asked to make an “old/new” discrimination for each trial. Scores ranged from 0 to 27 for old sequences and from 0 to 54 for new sequences.

We also assessed domain-general WM abilities with the Digit Span (DS) and Arithmetic subtests from the Working Memory index in Wechsler Adult Intelligence Scale IV ^41^ (WAIS – IV). During the DS subtest, participants listened to sequences of numbers of increasing length and were asked to repeat them in the same, inverse, or ascending order (DS Forward, DS Backward, and DS Sequencing, respectively). For the Arithmetic subtest, participants computed mathematical operations without external aids (e.g., paper and pencil, calculator). To compute individual WM abilities, we combined the raw scores from the DS and Arithmetic subtests. Scores ranged from 5 to 70.

Finally, formal musical training was assessed with the Goldsmiths Musical Sophistication Index ^42^ (Gold – MSI) questionnaire. This self-reported measure is comprised of 39 questions related to musical skills, experience, and habits. Each item is assessed on a 7-point Likert scale (from “1 = Completely disagree” to “7 = Completely agree”). Here, we used the Musical Training facet, which estimates an individual’s history of formal musical training. Individual scores on the Musical Training facet range from 7 to 49. Here, we used a Danish version of the Gold – MSI.

### Statistical analyses

Descriptive statistics were estimated for all variables. A one-way multivariate analysis of covariance (MANCOVA, Wilk’s Lambda [Λ], α = .05) was performed to compare STM, LTM, and WM skills between the two age groups while controlling for individual musical training level. We used one independent variable with two groups (young and elderly), five dependent variables (STM melody, STM rhythm, LTM old, LTM new, and WM scores) and one covariate (musical training score). The effect size was calculated using partial eta squared (i.e., partial η^2^).

Afterwards, to determine the effects of the independent variable and covariate, ten univariate analyses of covariance (ANCOVA) were computed individually for each of the dependent variables. These were computed at α = .005 after applying the Bonferroni correction (.05 divided by the number of ANCOVAs conducted) as follow-up tests to the MANCOVA.

## Results

### Descriptive statistics

Descriptive statistics for the measures of STM (melody and rhythm), LTM (old and new), WM, and musical training are provided in **Table 1**. Two outliers were removed from the old and new variables (z-scores below −3) and four values were missing from the WM measure. **Figure 1** illustrates group differences in normalized scores using raincloud plots ^43^.

**Table 1.** Descriptive statistics for short-term memory (STM melody and STM rhythm), long-term memory (LTM old and LTM new), working memory (WM), and musical training scores as a function of group. Mean and standard deviation scores are reported.

| Measure | Group |  |
| --- | --- | --- |
|  | Young (*N* = 37) | Elderly (*N* = 40) |
| STM melody | 14.22 ± 3.02 | 13.00 ± 3.22 |
| STM rhythm | 14.35 ± 2.65 | 12.53 ± 2.5 |
| LTM old | 23.14 ± 3.56 | 23.5 ± 4.47 |
| LTM new | 49.19 ± 5.2 | 43.21 ± 9.25 |
| WM | 43.00 ± 7.15 | 38.26 ± 7.67 |
| Musical training | 24.30 ± 6.88 | 22.18 ± 7.36 |

**Figure 1.**
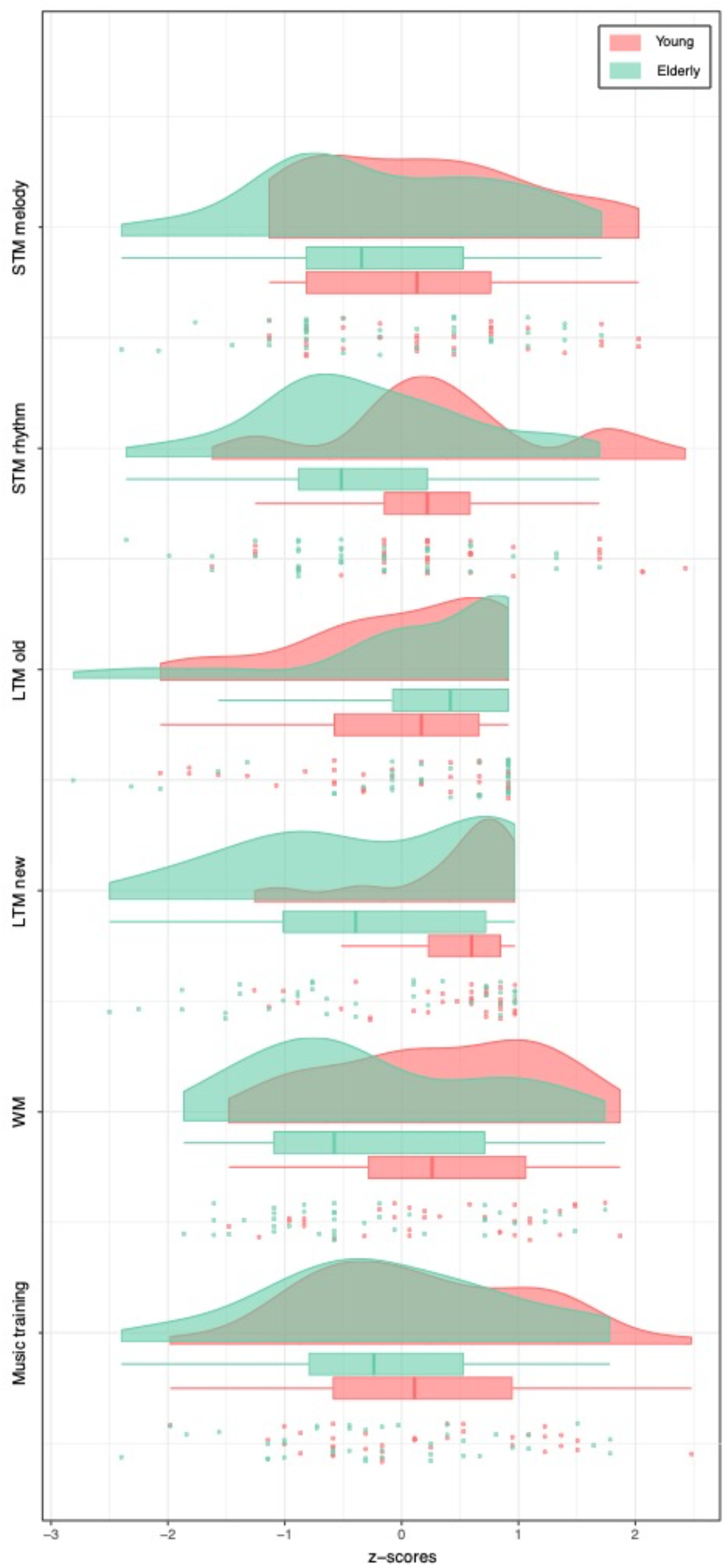
Raincloud plots show the overlapping distributions and normalized data points of both age groups in short-term memory (STM melody and STM rhythm), long-term memory (LTM old and LTM new), working memory (WM), and musical training. Boxplots show the median and interquartile (IQR, 25 – 75%) range, whiskers depict the 1.5*IQR from the quartile.

### Age-related differences in WM, STM, and LTM abilities

In order to assess the effect of age on WM, STM and LTM abilities whilst adjusting for musical training level, a one-way MANCOVA was used. Before computing the MANCOVA, we confirmed there were no significant differences in musical training level between the two groups (t(75) = 1.307, *p* = .195).

There was a statistically significant difference between the two groups on the combined dependent variables after controlling for musical training level (*F*(5, 64) = 4.129, *p* = .002, Wilks’ Λ = .756, partial η^2^ = .24). Follow-up ANCOVA showed statistically significant differences between young and elderly participants in STM rhythm (*F*(1, 74) = 10.299, *p* = .001, partial η^2^ = .12) and LTM new scores (*F*(1, 72) = 13.82, *p* < .001, partial η^2^ = .16). In addition, WM approached the significance level after correction for multiple comparisons (*F*(1, 70) = 7.631, *p* = .007, partial η^2^ = .1). On the contrary, STM melody (*F*(1, 74) = 3.535, *p* = .064, partial η^2^ = .05) and LTM old scores (*F*(1, 72) = 0.154, *p* = .696, partial η^2^ < .001) were not significant after controlling for musical training level.

There was a statistically significant effect of the covariate on the combined dependent variables (*F*(5, 64) = 3.858, *p* = .004, Wilks’ Λ = .768, partial η^2^ = .23). Follow-up ANCOVA showed statistically significant effects of musical training level on STM melody scores (*F*(1, 74) = 17.056, *p* < .001, partial η^2^ = .19) and LTM new scores (*F*(1, 68) = 13.46, *p* < .001, partial η^2^ = .16), meaning that higher level of musical training was associated to higher STM melody and LTM new scores. Musical training had no effect on WM (*F*(1, 70) = 3.285, *p* = .074, partial η^2^ = .04), STM rhythm (*F*(1, 74) = 5.744, *p* = .019, partial η^2^ = .07) and LTM old scores (*F*(1, 72) = 1.843, *p* = .17, partial η^2^ = .02).

## Discussion

The main goal of this study was to assess the effects of age and musical training on memory functioning. Our sample consisted of two age groups with matching musical training level. We found significant effects of age on specific memory abilities (i.e., STM for rhythm and LTM for new sequences). Additionally, we observed a positive effect of musical training level on certain auditory memory tasks.

Concerning STM capacity, we employed a musical paradigm that comprised same/different judgements of melodic and rhythmic musical sequences and confirmed that STM declines with age ^21, 22^. We found that young participants outperformed elderly participants in the STM rhythm measure, but there were no significant differences in the STM melody measure. These results are consistent with previous research on neural specializations for rhythm and melody. Using positron emission tomography (PET), Jerde et al. ^44^ showed that working memory for rhythm and melody activated distinct brain networks: the cerebellum, insular cortex, and cingulate gyrus for rhythmic sequences, and right frontal, parietal and temporal cortices for melodic sequences. Processing of rhythm has been typically linked to motor brain regions, such as the supplementary motor area, basal ganglia, and cerebellum ^45^. One explanation for the decreased performance we observed in the STM rhythm measure is the changes in cerebellar structure that occur with advancing age ^46, 47^. Indeed, one review linked age-related differences in cerebellar volume with cognitive and motor declines ^48^. This distinction between rhythm and melody was also evident when examining the effect of the musical training covariate. We found that musical training level had a positive effect on STM for melodic sequences, but not on rhythmic sequences. Since the musical training level of both groups was the same, it is reasonable that their performance was not significantly different in the STM melody measure.

To investigate LTM capacity, we asked participants to perform a musical recognition task. Once again, results differed depending on the variable analyzed: whereas young individuals outperformed older individuals in identifying musical sequences that were not presented before (“new”), the scores did not significantly differ when recognizing musical sequences that were previously listened to (“old”). Consistent with our results, older adults have been shown perform worse than young adults during free recall than recognition tasks ^49^. Moreover, elderly individuals exhibit increases in the number of false recognitions (i.e., remembering an event that did not happen) ^50, 51^. In terms of musical training level, we found a significant positive effect of this variable on the identification of new sequences, but no effect on the recognition of old sequences. This result denotes that memory recognition for musical stimuli was unaffected by previous musical experience. However, the distinction of old and new musical sequences required some musical skills.

We found no significant difference between the two age groups in WM skills. However, the results from the univariate ANOVA approached the significance level after correction for multiple comparisons, suggesting that young adults were close to outperform older adults. This is in line with previous studies that investigated age-related decline in WM abilities ^20, 52, 53^. In this study, we employed two WM tasks that are part of the widely used WAIS – IV. Performance on these tasks was previously shown to decline with age ^52-54^ and is accompanied by over- or under-activation of the dorsolateral prefrontal cortex ^25, 31, 32^. Such neural activity is coherent with the compensation-related utilization of neural circuits hypothesis (CRUNCH) ^55^, which suggests that over-recruitment of frontal and bilateral brain regions is a compensatory mechanism in older adults when task demands are low. However, as task load increases, neural resources are depleted, leading to neural under-recruitment and performance decline ^56^.

In addition to investigating differences in WM between both age groups, we also examined the effect of musical training on WM ability. We found no effect of the covariate on this measure. This result was expected, since the WM measure employed in this study had no relation to music or musical abilities. However, previous studies have shown positive effects of musical training and expertise on cognitive functioning ^36, 37, 57^ as well as relationships between musical training and preferences, cognitive abilities, and neuroplasticity ^58-65^, so it would be interesting to investigate this effect with the current WM measure in a sample of musicians and non-musicians.

Overall, our results indicate that (1) age has an effect on memory abilities, (2) not all memory types are equally affected by aging, and (3) musical training level has a positive effect on specific memory and musical skills. Furthermore, we found specific effects of aging on STM capacity for rhythmic sequences, but not for melodic sequences, and on LTM ability for identifying new sequences, but not for recognizing old sequences. Following previous studies on the spatiotemporal dynamics of memory ^66-69^, our future research will focus on correlating these differences in memory capacity to the patterns of brain activity and connectivity during short-term auditory memory recognition.

## Acknowledgements

We thank the Society for Education and Music Psychology (SEMPRE) for granting LB the SEMPRE’s 50th Anniversary Awards Scheme which was fundamental for realising this study.

The Center for Music in the Brain (MIB) is funded by the Danish National Research Foundation (project number DNRF117).

LB is supported by Carlsberg Foundation (CF20-0239), Center for Music in the Brain, Linacre College of the University of Oxford, and Society for Education and Music Psychology (SEMPRE’s 50th Anniversary Awards Scheme).

MLK is supported by Center for Music in the Brain, funded by the Danish National Research Foundation (DNRF117), and Centre for Eudaimonia and Human Flourishing funded by the Pettit and Carlsberg Foundations.

We thank Francesco De Benedetto for his assistance in the data collection.

## Data availability

The anonymized data from the experiment will be made available upon reasonable request.

## Author contributions

LB and EB conceived the hypotheses and designed the study. GFR, FC, ERO, OM, MK, and LB collected the experimental data. GFR and LB performed statistical analyses. GFR wrote the first draft of the manuscript. GFR and LB prepared the figures. PV, MLK, EB, and LB secured the fundings. MLK, PV, EB, and LB provided essential help to interpret and frame the results within the psychological and neuroscientific literature. All authors contributed to and approved the final version of the manuscript.

